# Reduced gap junction coupling amplifies the effects of cardiomyocyte variability and destabilizes the heartbeat

**DOI:** 10.1101/2025.03.20.644360

**Authors:** Karoline Horgmo Jæger, William E. Louch, Aslak Tveito

## Abstract

Cardiomyocytes exhibit significant cell-to-cell variability due to differences in protein expression and post-translational modifications in both the cell membrane and the intracellular machinery. Resulting variability in action potential propagation and configuration have been proposed to promote arrhythmia. However, such effects may be suppressed by tight electrical coupling of cells in the healthy heart, but not during pathological conditions where gap junction function is impaired. To investigate this question, we employed a cell-based mathematical model of cardiac electrophysiology, in which we systematically modified both the properties of individual cells within the array, and inter-cellular electrical connectivity (gap junctions). Despite the inclusion of marked variation in properties between cells, we observed electrical homogeneity across the array when cells were well coupled. In contrast, lower and/or more variable gap junction connectivity resulted in nonhomogeneous action potential configuration, and irregular timing of both the depolarizing and repolarizing electrical wavefronts. Pro-arrhythmic early after-depolarizations also occurred under these conditions, linked to reopening of L-type calcium channels. These effects were effectively dampened in highly coupled cells. Nevertheless, baseline differences in calcium homeostasis were not negated by gap junction coupling, indicating a limit to which electrical connections can homogenize mechanical function. There are also physical limits to electrical convergence, as we observed that action potential differences persisted at the edges and corners of the array where there are fewer electrical contacts with neighbouring cells. This finding may have implications for arrhythmic susceptibility in the border zone neighbouring an infarction. In summary, our findings underscore the critical role of intercellular coupling in maintaining cardiac stability and highlight the importance of studying cardiomyocytes within a syncytium rather than in isolation.

## 1 Introduction

The stability of the heartbeat depends on the coordinated electrical activity of *∼*2 billion cardiomyocytes. At the organ level, the healthy heart demonstrates remarkable electrical stability, even though individual cardiomyocytes exhibit significant variability in their electrophysiological and mechanical properties. Recent studies suggest that this variability is not merely tolerated but may play a vital role in enabling the heart’s adaptability and resilience, [1]. However, pathological changes that amplify or distort this variability can disrupt the delicate balance underpinning each heartbeat, leading to electrical instability, arrhythmogenesis, and impaired contractility. One critical factor in this context is the repolarization reserve, which represents the myocardium’s capacity to maintain stable repolarization despite variations in ion channel activity, [2]. When this reserve is compromised, electrical heterogeneities are magnified, increasing the likelihood of early afterdepolarizations (EADs) and arrhythmias, [3]. Fibrosis is another critical factor, as it introduces structural and electrical disruptions that amplify variability and reduce repolarization reserve. Specifically, fibrosis reduces gap junction coupling, slows conduction, and increases spatial heterogeneity, all of which may enhance arrhythmogenic potential, [4].

Several of these pathological alterations may be manifested during heart failure. This condition is commonly associated with myocardial fibrosis. In failing myocytes, electrophysiological remodeling can lead to a reduction in repolarizing potassium currents (*I*_Kr_ [5] and *I*_Ks_ [6]), which prolongs action potential duration (APD) and compromises repolarization reserve, further destabilizing electrical activity, [7, 8]. These effects promote reentrant circuits, EADs, and ventricular arrhythmias [4, 8]. Additionally, depolarizing currents such as late sodium current (*I*_NaL_ [5, 9]) and L-type calcium current (*I*_CaL_, [10, 11]) are upregulated, further increasing the likelihood of afterdepolarizations. These ion channel changes collectively impair electrical conduction and contribute to the pro-arrhythmic substrate in heart failure-related fibrosis.

In the present paper, we use recently developed mathematical models to demonstrate the strong effect of cell-to-cell electrical coupling on the action potential of individual cardiomyocytes and its influence on the stability of the excitation front. Even with substantial cell-to-cell variations in membrane properties, a stable excitation front is maintained if the gap junction coupling is sufficiently strong. Conversely, when coupling is reduced, the excitation front becomes irregular, creating a substrate for arrhythmias.

The findings described above suggest that arrhythmic susceptibility during pathology is linked to cardiomyocyte electrical remodeling, as well as increased between-cell variability in electrophysiology. While well-connected cardiomyocytes might be expected to negate such electrical differences, we hypothesized that impaired cellular connectivity during disease may compromise this capacity, and provide a substrate for arrhythmia.

Mathematical models of cardiac electrophysiology fall into two main categories: one focuses on modeling the action potentials and calcium dynamics of single cardiomyocytes, [12, 13, 14], while the other focuses on modeling cardiac tissue to study spatial dynamics, [15, 16, 17, 18, 19]. Single cell models excel at detailing intracellular processes but lack the tissue-scale resolution needed to capture phenomena such as reentry and fibrillation. Importantly, cardiomyocytes behave significantly differently as isolated units compared to when they are electrically coupled within a syncytium, [20]. Spatial tissue models address the vast computational problem posed by huge numbers of cardiomyocytes by homogenizing tissue properties, enabling simulations of large-scale conduction and arrhythmias, see, e.g., [15, 21, 22].

Traditionally, spatially resolved mathematical models of cardiac tissue, including the widely used bidomain model and the simpler monodomain model, have assumed that the extracellular (E) space, cell membrane (M), and intracellular (I) space overlap and are present everywhere. While this assumption is fundamentally incorrect, these models have been extensively and successfully used to explain key phenomena in cardiac electrophysiology. However, when studying the effects of local cell-to-cell variations in membrane properties and heterogeneity in gap junction coupling, homogenized models fall short. This is because individual cells and gap junctions are not explicitly represented in the mathematical formulation of these models.

Recent efforts have focused on explicitly representing the E, M, and I spaces of cardiomyocytes in mathematical models, [23, 24, 25]. The resulting EMI models significantly improve accuracy by allowing ion channel densities to vary along the cell membranes, [26]. However, this improvement comes at the cost of a substantial increase in computational complexity, [27]. To address this, an intermediate model, referred to as the Kirchhoff network model (KNM, [28]), has been developed. The KNM enables the representation of individual cells with distinct properties, and distinct cell-to-cell coupling, but does not allow for variations in membrane densities along individual cells.

As the bidomain model has been successfully approximated by the monodomain model, the KNM can be further simplified into the SKNM (simplified KNM), [29]. In [30], the computational efficiency of these models was evaluated by simulating 4000 timesteps for a domain of 13*×*65 physical cells. The results showed the following CPU times: 3 seconds for SKNM, 7 seconds for KNM, 16 hours for the EMI model, 28 minutes for the bidomain model, and 4 minutes for the monodomain model. Both the KNM and SKNM provided results very close to those of the EMI model, which remains the most accurate approach.

Using the cell-based SKNM, we, in this study, investigate how the action potentials of individual cardiomyocytes in tissue are influenced by reduced gap junction coupling. We demonstrate that weakened coupling leads to a pronounced divergence in the action potential across cells in a tissue with randomized properties. This divergence is amplified at the boundaries of the excitable region, where cells experience both reduced coupling and the effects of being at the tissue edge. These findings have implications for the border zone of infarcted regions, where gap junction coupling is commonly diminished, and boundary effects further amplify electrical heterogeneities, see, e.g., [31].

Furthermore, we show that reduced gap junction coupling destabilizes excitation fronts, making them irregular, and causes the repolarization wave to become ragged. These results highlight the intricate interplay between gap junction coupling, cellular heterogeneity, and tissue boundaries in maintaining electrical stability.

A key consequence of disrupted electrical coupling in heterogeneous cardiac tissue is the possible formation of EADs. These secondary depolarizations can arise when repolarization reserve is diminished, leading to prolonged action potentials and destabilized membrane dynamics. We demonstrate that while EADs do not emerge under normal coupling conditions, the combination of reduced gap junction conductance and increased cell-to-cell variability creates a vulnerable substrate where EADs can arise. This suggests that electrical heterogeneity and impaired coupling may jointly contribute to proarrhythmic conditions, further emphasizing the critical role of intercellular communication in maintaining cardiac stability.

## 2 Methods

### 2.1 Cell-based model of cell collections — The simplified Kirchhoff network model (SKNM)

We simulate small collections of cardiomyocytes using the simplified Kirchhoff network model (SKNM), [29]. In this model, cells are electrically coupled to neighboring cells, and the membrane and intracellular dynamics of each cell are represented. For each cell, *k*, the model reads

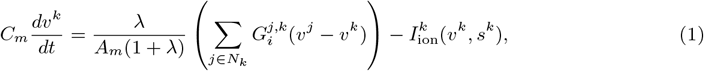

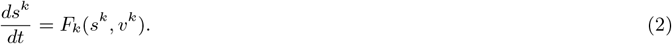

Here, *v*^*k*^ is the membrane potential of cell *k* (in mV), *N*_*k*_ is a collection of all the neighboring cells of cell 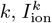 is the sum of the transmembrane current densities of cell *k* (in A/F), *s*^*k*^ are additional cell variables including intracellular concentrations and ion channel gating variables, and *F*_*k*_ governs the dynamics of these variables. Furthermore, *C*_*m*_, *A*_*m*_, *λ* and 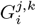 are model parameters specified in Section 2.1.1. The models used for the single cell membrane and intracellular dynamics, i.e., 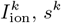 and *F*_*k*_, are described in Section 2.2.

#### 2.1.1 SKNM parameters

We consider a 2D grid of cells where each cell is connected to its direct neighbors in the *x*- and *y*-directions (i.e., no diagonal cell connections). Furthermore, we assume that the cells are shaped as cylinders with length *l* = 100 *µ*m and diameter *d* = 20 *µ*m.

In SKNM (1)–(2), *C*_*m*_ is the specific membrane capacitance, set to *C*_*m*_= 1 *µ*F/cm^2^ [32], and *A*_*m*_ (in cm^2^) is the membrane area, set to *A*_*m*_ = 2*πld*. Here, *πld* is the geometrical surface area of the cells (except for the cell ends, which are assumed to consist of cell connections). The factor 2 is included to account for the fact that the actual (capacitive) membrane area is larger than the geometrical membrane area due to membrane folding (see, e.g., [33, 34, 35, 36]). The parameter 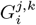 represents the intercellular conductance and is given by

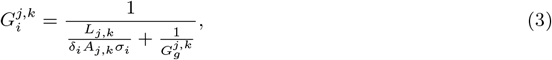

where *L*_*j,k*_ is the distance between the centers of cells *j* and *k* (in cm), *δ*_*i*_ is the intracellular volume fraction (set to 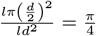), *A*_*j,k*_ is the cross sectional area between cells *j* and *k, σ*_*i*_ = 4 ms/cm is the intracellular conductivity [37, 38], and 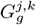 is the gap junction conductance between cells *j* and *k* (in mS).

The parameter *λ* is a unitless constant introduced to simplify KNM to SKNM (see [28, 29]), and should satisfy

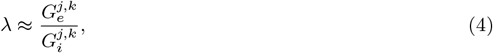

Where 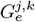 is the extracellular conductance, given by

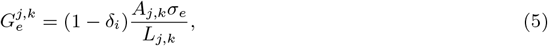

and *σ*_*e*_ = 20 mS/cm is the extracellular conductivity [39]. In our simulations, the value of 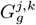 is often allowed to vary between cell connections, and *L*_*j,k*_ and *A*_*j,k*_ varies depending on whether the cells *j* and *k* are connected in the *x*- or *y*-directions. Therefore, there exists no constant *λ* such that 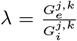 for all *j* and *k*, and we need to define *λ* such that the approximation (4) holds. To this end, we define *λ* by applying the average gap junction conductance, 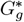, and by using the conductances, 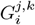 and 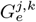, for cell connections in the *x*-direction. That is, we set

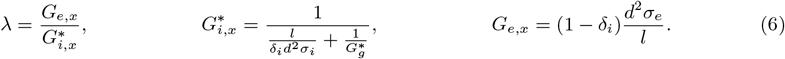

### 2.2 Models for single cell intracellular and membrane dynamics

In this section, we describe the models used for the single cell intracellular and membrane dynamics defined in 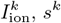, and *F*_*k*_ in (1) and (2) above.

#### 2.2.1 Human ventricular base model

For most of our simulations and unless otherwise stated, we use a version of the adult ventricular base model from [40] to model the intracellular and membrane dynamics.

To allow for investigations of contraction synchrony, we have incorporated a model for mechanical cell contraction in the base model for some of our simulations (Figures 3–8), as was also done in [41]. We represent cell contraction using the Rice et al. model [42]. This model represents cell contraction in the form of shortening of the sarcomere length. In order to obtain a reasonable sarcomere length shortening for the base model, the parameters *k*_on_ and *PCon*_titin_ in the Rice et al. model are set to 135 *µ*M^*−*1^s^*−*1^ and 0.018, respectively. In addition, the total concentration of cytosolic calcium buffers in the base model, 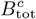, is set to 0.098 mM and half of this concentration is assumed to be troponin. Otherwise, the parameters of the original models from [40, 42] are applied.

#### 2.2.2 Early afterdepolarizations (EADs) — The O’Hara-Rudy model

To investigate properties of early afterdepolarizations (EADs) in heterogeneous cell collections, we apply the O’Hara-Rudy model [43] for the ventricular action potential, which exhibits EADs for extensive *I*_Kr_ block in combination with slow pacing. Because the O’Hara-Rudy model is known to exhibit un-physiologically slow conduction velocities (see, e.g., [44, 45]), we have replaced the *I*_Na_ model from the O’Hara-Rudy model with the version found in the ten Tusscher model [46], as suggested in a comment by the authors of the O’Hara-Rudy model [47]. In addition, the sodium channel conductance, *g*_Na_, of the ten Tusscher model was multiplied by a factor 2 to allow for a conduction velocity of about 50 cm/s in the normal healthy case.

**Table 1:** Overview of the different cases of variation in cell properties and gap junction coupling considered in our simulations.

|  | Cell variation | Gap junction variation | Variation distribution |
| --- | --- | --- | --- |
| Case 1 | Two cell types | No | Random |
| Case 2 | Two cell types | No | Systematic |
| Case 3 | None | Yes | Random |
| Case 4 | Cell parameters | No | Random |
| Case 5 | Cell parameters | Yes | Random |
| Case 6 | Perturbation currents | No | Random |

#### 2.2.3 Stimulation protocols

In all our simulations (except for the cases with no gap junction coupling), we initiate action potentials by stimulating the first (leftmost) row of cells in the *x*-direction. For the cases with no gap junction coupling, we stimulate all cells. For the base model and for the O’Hara-Rudy model with no drug present, we use 1 Hz pacing and for the O’Hara-Rudy model with *I*_Kr_ block we use 0.25 Hz pacing. To allow the model variables (e.g., ion concentrations) time to adjust to model parameter changes, we stimulate the cell collections 60 times for the base model and 200 times for the O’Hara-Rudy model before recording the solutions. The difference between models is due to a slower convergence of variables in the O’Hara-Rudy model compared to the base model (cf., e.g., [48]).

### 2.3 Variation of cell and gap junction properties

To investigate the effect of heterogeneous cell properties, we consider cell collections with variations in the properties of the cells and in the strength of the gap junction coupling between cells.

#### 2.3.1 Variation of cell properties

In this subsection we describe the considered variations in single cell properties. The specific cases of variation applied in our simulations are summarized in Table 1.

##### Two cell types case

As a first example of cell collections with varying cell properties, we consider the mix of just two different types of cells. In this case, one type of cell is referred to as healthy and is characterized by the default ventricular base model described in Section 2.2.1. The other type of cell is also modeled using the ventricular base model, but for these cells the duration of the action potential is considerably increased by setting the *I*_Kr_ current to zero, reducing the conductance of the *I*_K1_ current by 75%, and doubling the conductance of the *I*_CaL_ current.

##### Random variation in cell parameters

The second type of considered cell variation is a random variation in cell properties between each cell. This is represented by randomly varying the parameters representing the maximum conductance of ion channels and the maximum currents of pumps and exchangers in the cell membrane and for intracellular calcium fluxes. In addition, the total concentration of different calcium binding buffers is allowed to vary. More specifically. we vary the value of the parameters 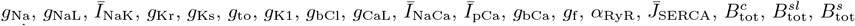, and 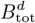 in the base model, and the parameters 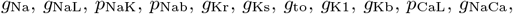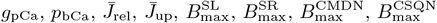, and 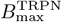 in the O’Hara-Rudy model.

To enable investigation of different degrees of cell variation, we draw a random number *α*_*i,k*_ between *−*1 and 1 for each cell *k* and each parameter *i*. The parameter *p*_*i*_ is then redefined for each cell *k* as

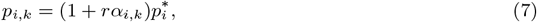

Where 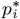 is the default parameter value in the original single cell model and *r* is a unitless scaling parameter specifying the degree of variation for the considered cell collection. For instance, *r* = 0.5 is referred to as 50% cell variation.

##### Constant perturbation currents

As a third example of varying cell properties, we define a set of perturbation currents. These are constant in time, but the constant value varies between the cells in the cell collection. Again, to investigate the effect of increasing the degree of cell variation, the perturbation currents are defined on the form

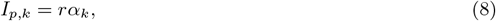

where *I*_*p,k*_ is the perturbation current for cell *k*, in (A/F), *r* is a unitless scaling parameter specifying the degree of variation for the considered cell collection, and *α*_*k*_ is a value between *−*1 A/F and 1 A/F drawn randomly for each cell. The constant perturbation current is incorporated in the model by redefining 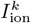 in (1) to

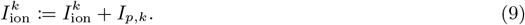

#### 2.3.2 Variation of gap junction coupling

In addition to varying the properties of the individual cells, we also consider random variations in the gap junction coupling between cells. For each cell connection, *i*, we draw a random number *β*_*i*_ and set up the gap junction coupling for cell connection *i* to be

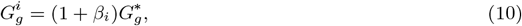

Where 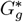 is the average gap junction conductance in the cell collection. The value of *β*_*i*_ is drawn randomly from the interval *−*0.25 to 0.25 for 25% gap junction variation and from the interval *−*0.5 to 0.5 for 50% gap junction variation. We also consider cases with a constant gap junction coupling (0% variation). In that case, 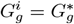 for all cell connections *i*.

In our investigations, we consider several different values of the average gap junction coupling, 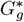. For normally connected cells with a longitudinal conduction velocity of about 50 cm/s, the value of 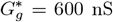 is applied. In addition, we consider several cases of reduced gap junction coupling. The weakest considered coupling strength is 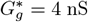 for cells modeled using the base model and 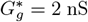 for cells modeled using the O’Hara-Rudy model. These are the smallest values of 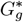 which allow for a traveling excitation wave for the two considered models. An exception is for the base model case with a smix of healthy and perturbed cells (Cases 1 and 2 in Table 1). For the perturbed cells, the conductance of the *I*_K1_ current is reduced, which makes the cells more excitable. For this case, we observe that a coupling strength of 2 nS is sufficient to allow for an excitation wave, even for collections of a mix of normal and perturbed cells (see Figures 1 and 2).

**Figure 1:**
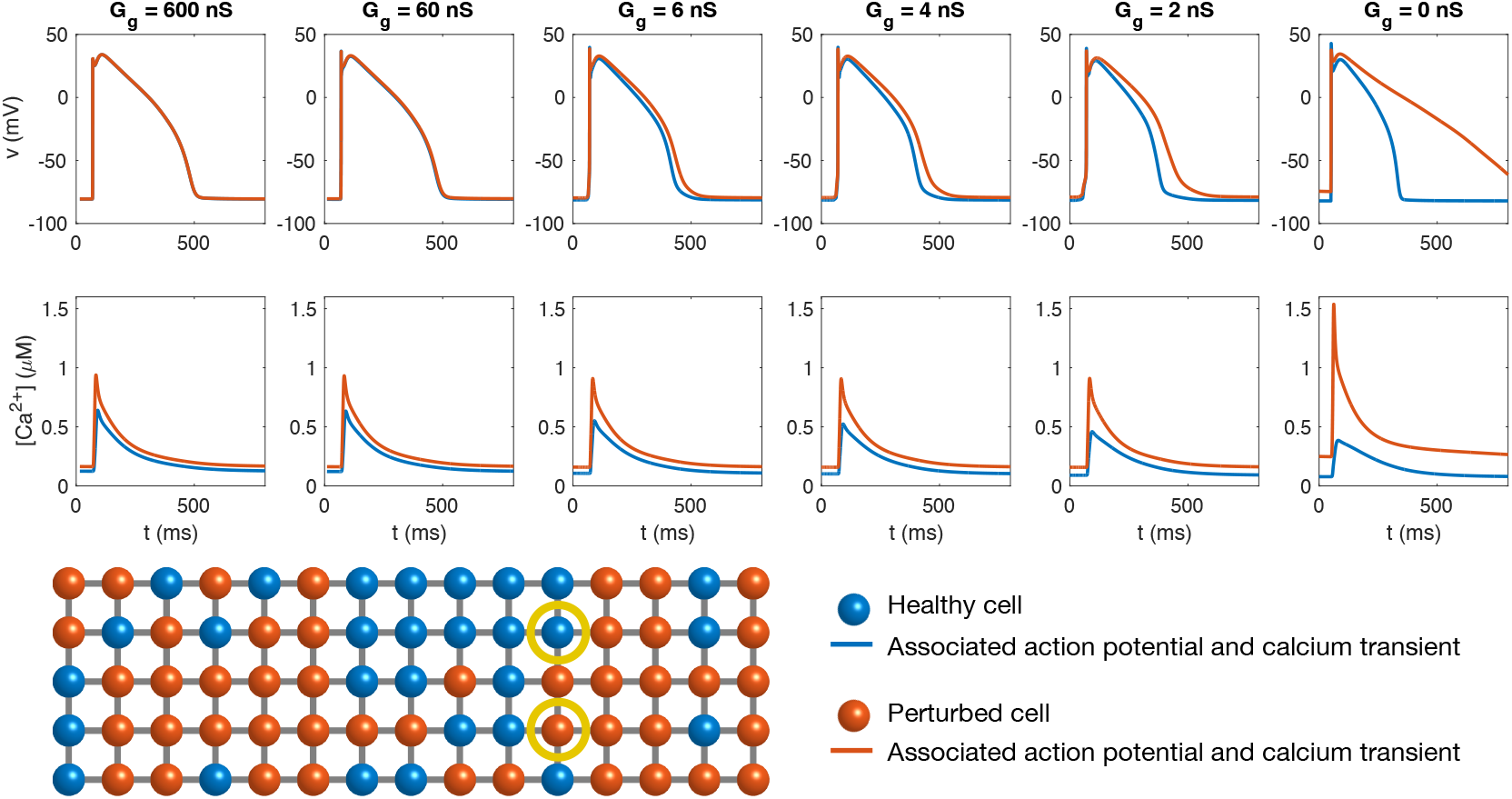
Action potentials and calcium transients in simulations of 15*×*5 randomly distributed cells of two different types (blue: normal, healthy cells, red: cells with no *I*_Kr_, double *g*_CaL_, and 75% reduced *g*_K1_), Case 1 in Table 1. In the leftmost panel, the cells are strongly connected (600 nS) like in normal healthy tissue. The cell connections are gradually reduced in the right panels, as indicated in the panel titles. In the case with *G*_*g*_ = 0 nS, the cells are completely disconnected. The lower panel illustrates the distribution of cell types in the cell collection, and the yellow circles mark the cells whose action potentials and calcium transients are reported in the upper panels. The gap junction coupling is the same between all cells in the collection.

**Figure 2:**
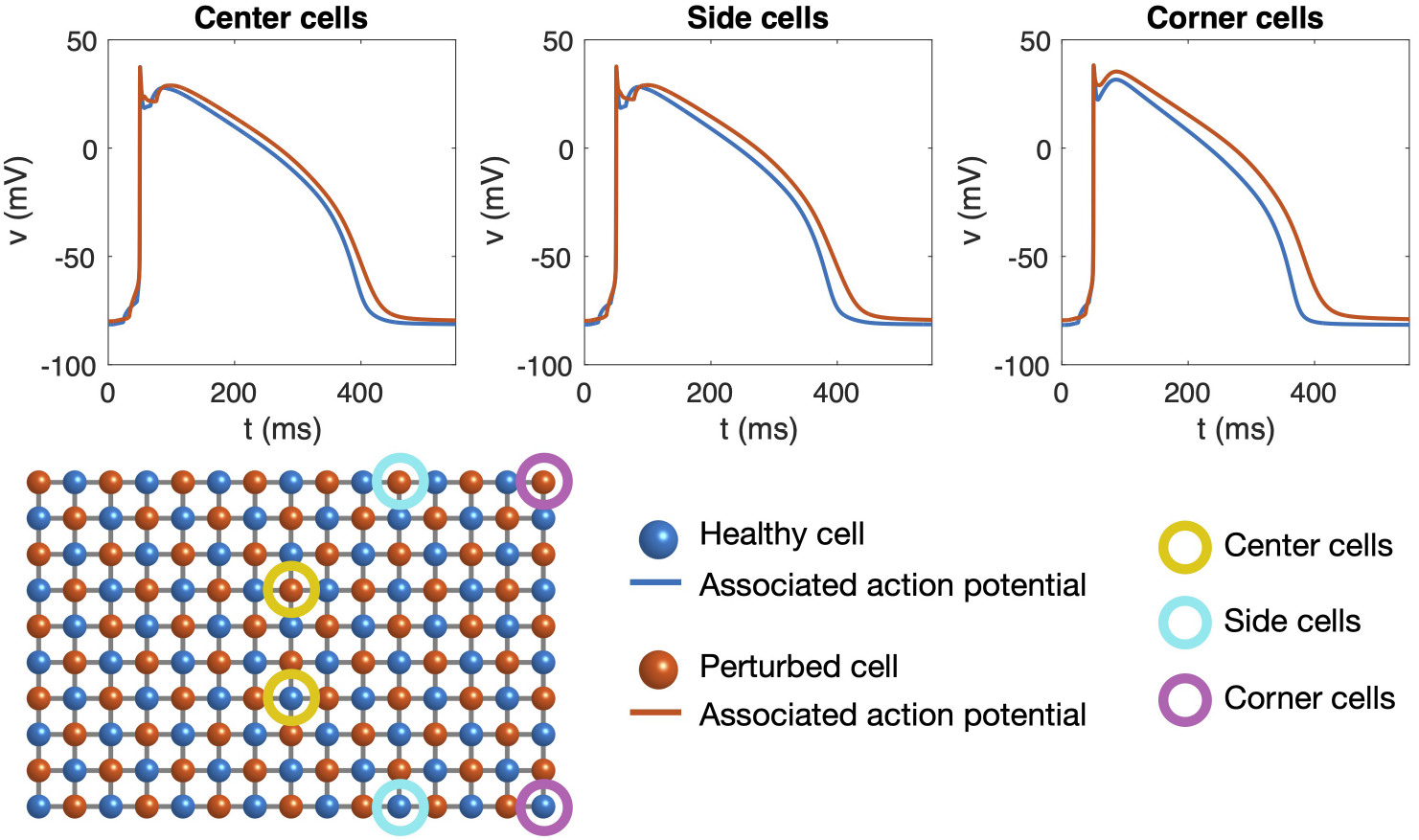
Action potentials recorded in six different cells in a simulation of 15*×*20 cells. The cell collection consists of two different types of cell (blue: normal, healthy cells, red: cells with no *I*_Kr_, double *g*_CaL_, and 75% reduced *g*_K1_), Case 2 in Table 1. The cell types are evenly distributed throughout the domain such that every other cell is normal and every other cell is perturbed (see lower panel). The gap junction conductance is set to 2 nS for all cell connections. In the upper left panel, we compare the action potential of the two cell types for cells located in the center of the cell collection (marked with yellow circles in the lower panel). In the upper center panel, we compare the action potential of the two cell types for cells located at sides of the cell collection (marked with turquoise circles in the lower panel). Finally, in the upper right panel, we compare the action potential of the two cell types for cells located in corners of the cell collection (marked with purple circles in the lower panel).

**Figure 3:**
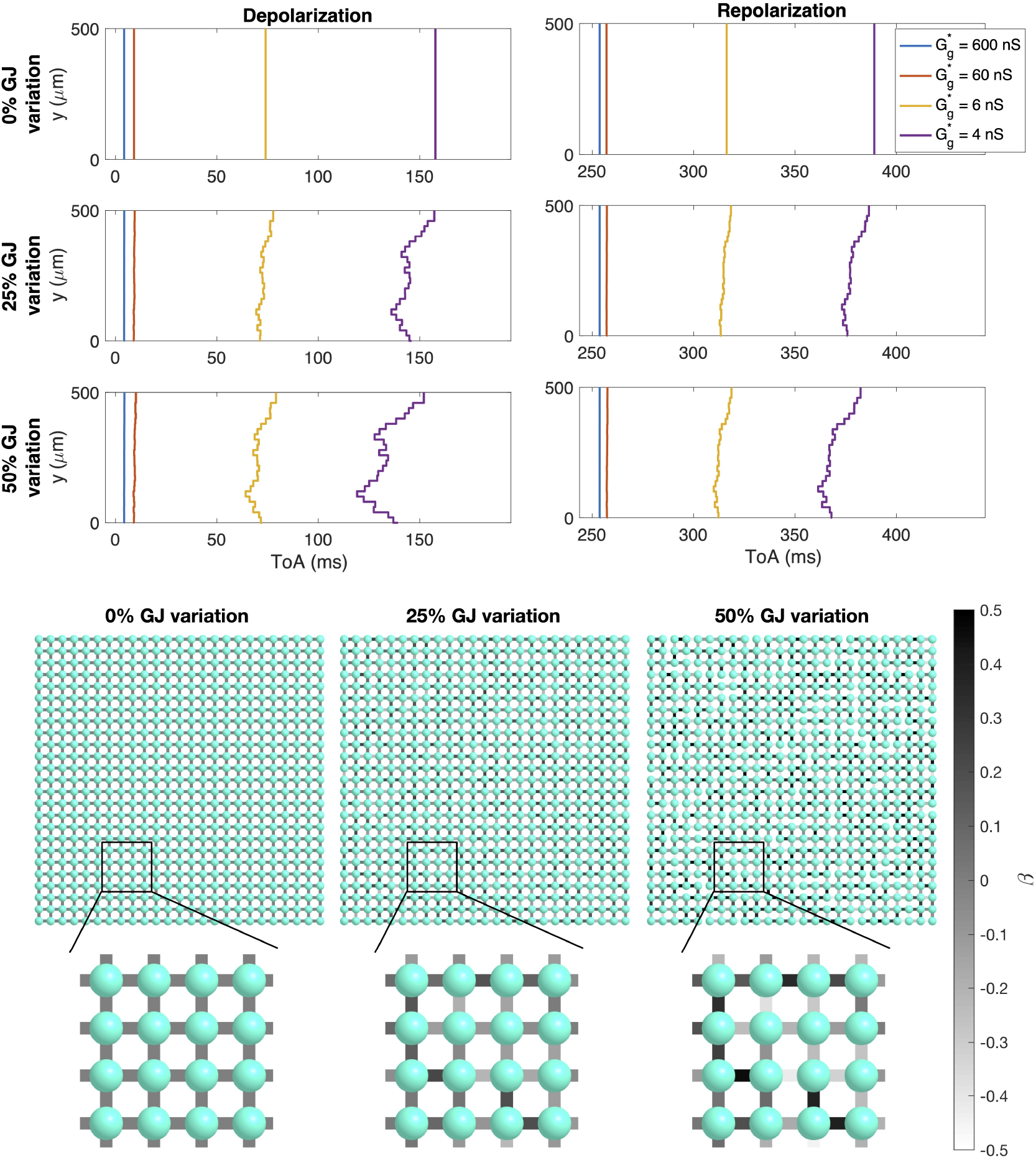
Simulations of 25*×* 25 cells with different degrees of gap junction coupling variations and different average gap junction conductances (Case 3 in Table 1). Upper panel: Time of arrival (ToA) for the depolarization and repolarization waves for cell number 10 in the *x*-direction and each of the cells in the *y*-direction. The cell properties are the same for all cells, but the degree of gap junction coupling variation increases for each plot row. The different lines correspond to different values of the average gap junction conductance, 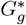 (see legends). Note that the ToA axis covers different time points in the different plots, but the scaling of the axis is the same in all cases. Lower panel: Illustration of cell collections with 0%, 25% and 50% variations in the gap junction coupling. The color of the cell connections indicate the strength of the gap junction coupling between cells, *β*, in the sense that 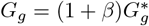.

### 2.4 Definition of biomarkers

In this section, we define the biomarkers considered in our investigations.

#### 2.4.1 Action potential duration (APD)

In order to illustrate the difference in cell properties resulting from random variation of cell properties, we display the action potential duration for the different cells computed for a case when the cells are not connected (see, e.g., the lower panel of Figure 4). We display the APD50 value defined as the duration from the time of the maximum upstroke velocity until the time when the membrane potential crosses halfway between the maximum and minimum membrane potential values during repolarization.

**Figure 4:**
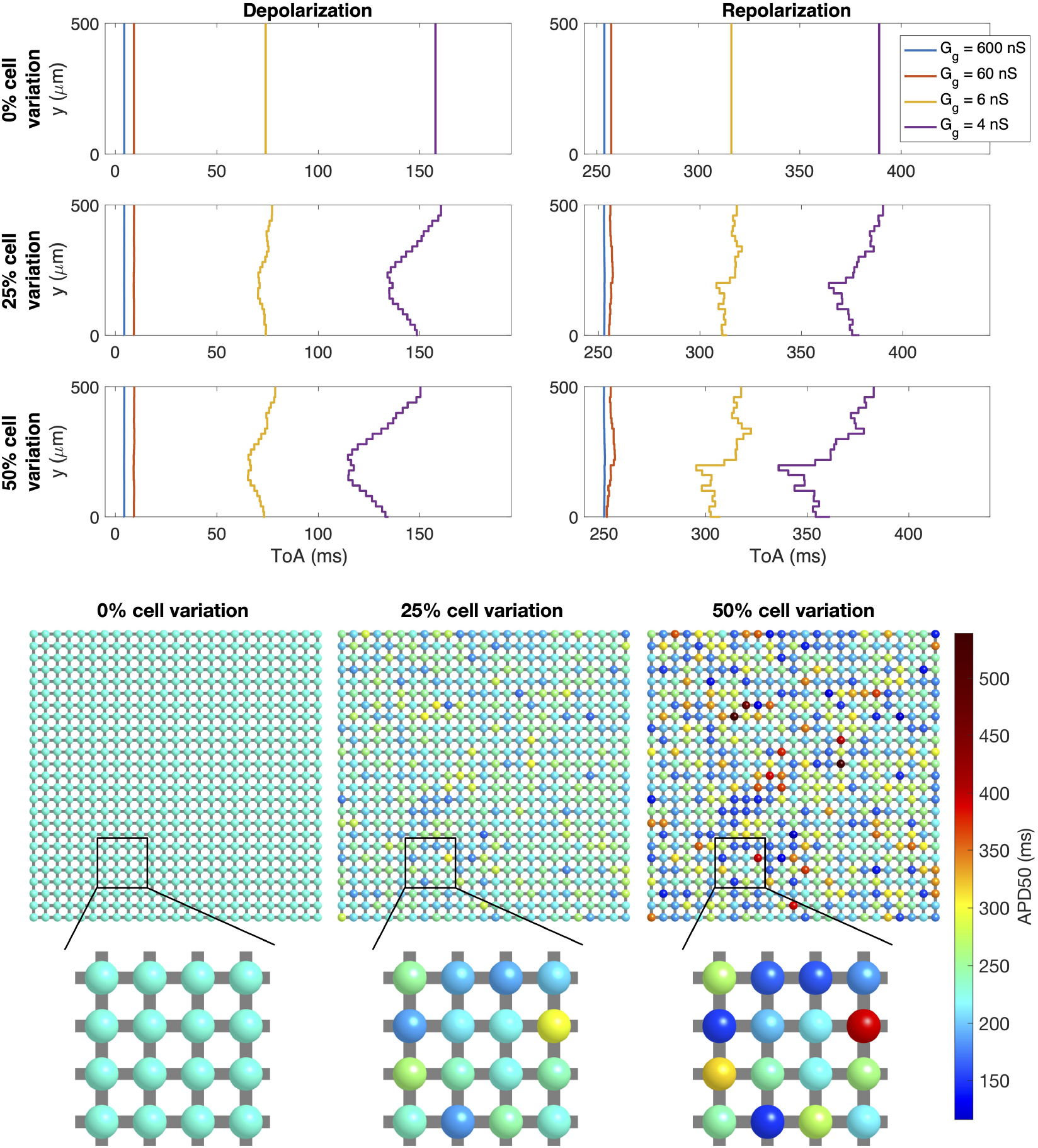
Simulations of 25 *×* 25 cells with different degrees of cell variations and different average gap junction conductances (Case 4 in Table 1). Upper panel: Time of arrival (ToA) for the depolarization and repolarization waves for cell number 10 in the *x*-direction and each of the cells in the *y*-direction. The degree of variation of cell properties are increased for each plot row, but the gap junction conductance is the same for all cell connections. The different lines correspond to different values of the average gap junction conductance, 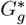 (see legends). Note that the ToA axis covers different time points in the different plots, but the scaling of the axis is the same in all cases. Lower panel: Illustration of the cell collections with 0%, 25% and 50% variations in the cell properties. The color of the cells indicate the APD50 value of the cell if it was not connected to other cells. Note, however, that the variation in cell parameters affect several different cell properties, not only the APD50 value.

#### 2.4.2 Time of arrival (ToA) during depolarization and repolarization

To investigate the shape of the wavefront during cellular depolarization and repolarization, we define the time of arrival (ToA) for a cell as the point in time when the membrane potential crosses *−*30 mV during depolarization or repolarization. We compare this ToA for cells located at the same distance from the stimulation location, i.e., for cells located at the same *x*-value, but for different *y*-values.

### 2.5 Numerical methods

In our computations, the model (1) and (2) is formulated as a coupled system of ordinary differential equations and solved using the *ode15s* function in MATLAB.

## 3 Results

### 3.1 Effects of cell heterogeneity is evened out in collections of connected cells

A first illustration of how effects of cell heterogeneity are evened out in collections of randomly-arranged, connected cells is given in Figure 1. In this figure, we consider a collection of two types of cells, one healthy and one perturbed case with an extra long action potential caused by a reduction in *I*_Kr_ and *I*_K1_ and an increase in *I*_CaL_ (Case 1 in Table 1). The action potentials of the two cell types in the case when there is no connection between the cells are displayed in the rightmost figure panel and exhibit a considerable difference in action potential duration. In the panels to the left, we similarly plot the action potentials of one cell of each type in simulations where the cells are electrically connected through gap junctions. In these cases, we observe that the differences between the two cell types are evened out, and for a high degree of cell coupling, the action potentials in the two cell types are completely indistinguishable. Note that APD50 is 227 ms and 461 ms for stand alone cells of the two types, whereas APD50 is 342 ms for both cell types when the cells in the collection are connected with the strongest considered gap junction coupling.

Figure 1 also displays the calcium transients of the two cell types. In the case with no cell coupling, the perturbed cell display a considerably higher calcium concentration level and a higher maximum amplitude of the calcium transient. For the coupled cases there is less of a difference in the calcium transients, as differences in cellular action potential are negated. As expected, calcium transients remain somewhat larger in the cell with the larger *I*_CaL_ current, even when this cell is highly coupled to its neighbors.

Comparing the panels of Figure 1 from left to right, we observe that for a normal degree of cell-to-cell coupling (*G*_*g*_ = 600 nS), the action potentials for the two cell types overlap completely, whereas for a considerable reduction in gap junction coupling (e.g., *G*_*g*_ = 2 nS), differences between the two cell types’ action potentials are visible. This shows that the effects of cell heterogeneity is more pronounced for reduced cell coupling. This property will be further illustrated in other examples below.

### 3.2 Cell heterogeneity is more pronounced at cell collection boundaries

In Figure 2, we consider a case of evenly distributed cells of the two cell types considered in Figure 1 (Case 2 in Table 1). In this case, we consider weak (2 nS) gap junction coupling and compare the action potentials of one cell of each type when the cells are located in the center of the cell collection and when the cells are located at the sides or in the corners of the cell collection. We observe that the difference between the two cell types increases from the center cells to the side cells to the corner cells. More specifically, APD50 differs by 12 ms for the two cells in the center, and it differs by 17 ms for the side cells and by 28 ms for the corner cells. This difference between cell locations may be explained by the fact that the corner and side cells are coupled to fewer cells and thereby less affected by neighbouring cells.

### 3.3 Depolarization and repolarization wavefront irregularities are more pronounced for reduced cell coupling

#### Variation in gap junction coupling

In Figure 3, we consider simulations of 25*×*25 cells. Here, all cells have the same properties, but three different degrees of variation in the gap junction coupling (0%, 25%, and 50%) and four different values for the average gap junction conductance (600 nS, 60 nS, 6 nS, and 4 nS) are considered (Case 3 in Table 1). We plot the time of arrival (ToA) for the depolarization and repolarization waves along different values of *y* for the same value of *x*. In the upper panel, there is no variation in gap junction coupling and the ToA is the same for all the cells along the vertical line. The ToA varies between the different values of the average gap junction coupling in the sense that the wave reaches the considered *x*-value faster when the gap junction strength is high. In the next panels the degree of gap junction variation is increased to 25% and 50%, and we observe irregularities in the ToA in the cases of weak cell coupling, especially for the depolarization wave, but also to some extent for the repolarization wave. For a normal cell coupling 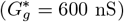, however, there appears to be no visible variation in the ToA, even in the cases of high gap junction coupling variation.

#### Variation in cell properties

In Figure 4, we consider variations in cell properties instead of variations in gap junction coupling (Case 4 in Table 1). The cell properties are set up by randomly varying model parameters like explained in Section 2.3.1. Some of the differences in cell characteristics caused by varying properties is illustrated in the lower panel of the figure, which displays the APD50 values of each cell if it was observed in isolation. We observe that for 25% cell variation, APD50 varies between about 170 ms and 300 ms, whereas for 50% variation the APD50 varies between about 120 ms and 540 ms. In the upper panel of Figure 4, similar tendencies as those displayed for the gap junction variation in Figure 3 are observed. For example, for no cell variation or strong gap junction coupling, the ToA curves are straight, but when cell variation is introduced, irregularities are present for a weak cell coupling. In this case, the maximum variation appears to be quite similar for the depolarization and repolarization waves, but there appears to be more raggedness for the repolarization wavefront.

#### Variation in cell properties and gap junction coupling

The effects of gap junction variation and cell variation are combined in the simulations reported in Figure 5 (Case 5 in Table 1). Here, we again observe straight wavefronts for normal cell coupling, but irregular wavefronts for a reduced cell coupling. The results observed in Figure 3–5 are summarized in Figure 6. In this figure, the maximum difference in ToA for the same value of *x* are plotted as functions of the average gap junction coupling, 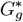. We observe that whether there is gap junction coupling variation or not, the maximum difference in ToA during depolarization and repolarization increases as the degree of cell variability is increased and as the gap junction coupling is reduced.

**Figure 5:**
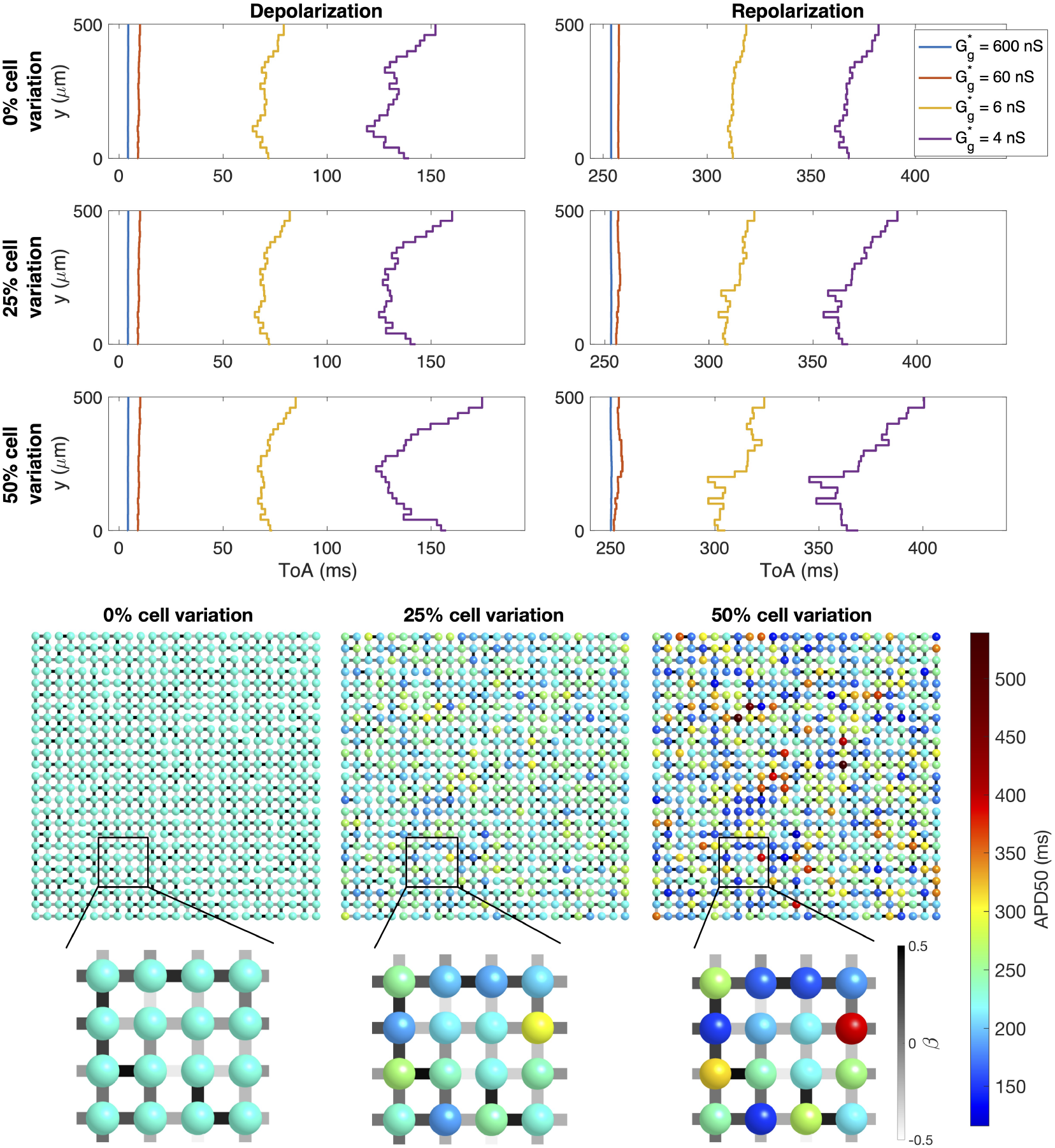
Simulations of 25 *×* 25 cells with 50% variation in gap junction coupling, different degrees of cell variations and different average gap junction conductances (Case 5 in Table 1). Upper panel: Time of arrival (ToA) for the depolarization and repolarization waves for cell number 10 in the *x*-direction and each of the cells in the *y*-direction. The degree of variation of cell properties are increased for each plot row. The different lines correspond to different values of the average gap junction conductance, 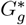 (see legends). Note that the ToA axis covers different time points in the different plots, but the scaling of the axis is the same in all cases. Lower panel: Illustration of the cell collections with 0%, 25% and 50% variations in the cell properties. The color of the cells indicate the APD50 value of the cell if it was not connected to other cells. Note, however, that the variation in cell parameters affect several different cell properties, not only the APD50 value. The color of the cell connections indicate the strength of the gap junction coupling between cells.

**Figure 6:**
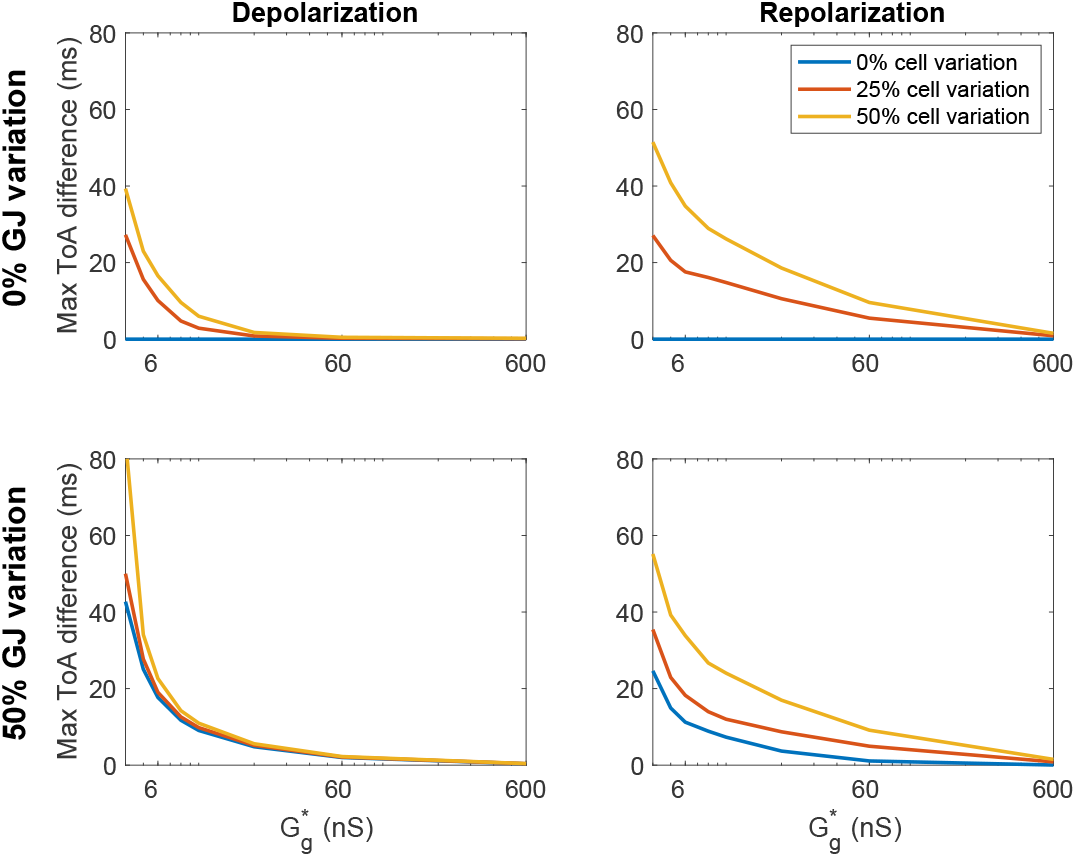
Maximum variation in the time of arrival (ToA) for the depolarization and repolarization waves as functions of the average gap junction conductance, 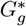, for the simulations described in Figures 3–5 (Cases 3–5 in Table 1).

### 3.4 Differences in calcium transients and cell contraction are not necessarily evened out for increased gap junction coupling

In Figure 7, we investigate properties of the simulations in Figures 3–6 further by plotting the action potential, calcium transient and the sarcomere length shortening of two selected cells in the cell collection. More specifically, we consider the two cells whose action potential duration differ most at the same *x*-value for the simulation with 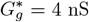. The parameter adjustments for these two cells are displayed in the lower panel of Figure 7. For instance, we observe that there are large differences in 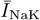 and *g*CaL between the two cells. In addition, 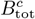 is considerably higher for the red cell than for the blue cell, which is likely the reason for the reduced calcium transient amplitude and sarcomere length shortening observed for the red cell compared to the blue cell. We consider the case of 50% variation in both gap junction coupling and cell properties (Case 5 in Table 1) and four different values of the average gap junction coupling, 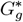. We observe that like in Figure 1, the difference in action potential shape decreases as the gap junction coupling strength is increased. On the other hand, the difference in calcium transient and sarcomere length shortening is present even for a strong gap junction coupling. In fact, for these example cells, the difference in calcium transient and contraction amplitude increases as the gap junction coupling increases. This is likely explained by the fact that for weaker cell coupling, the APD is shorter (and thereby the open time for *I*_CaL_ is also shorter) for the cell corresponding to the blue line.

**Figure 7:**
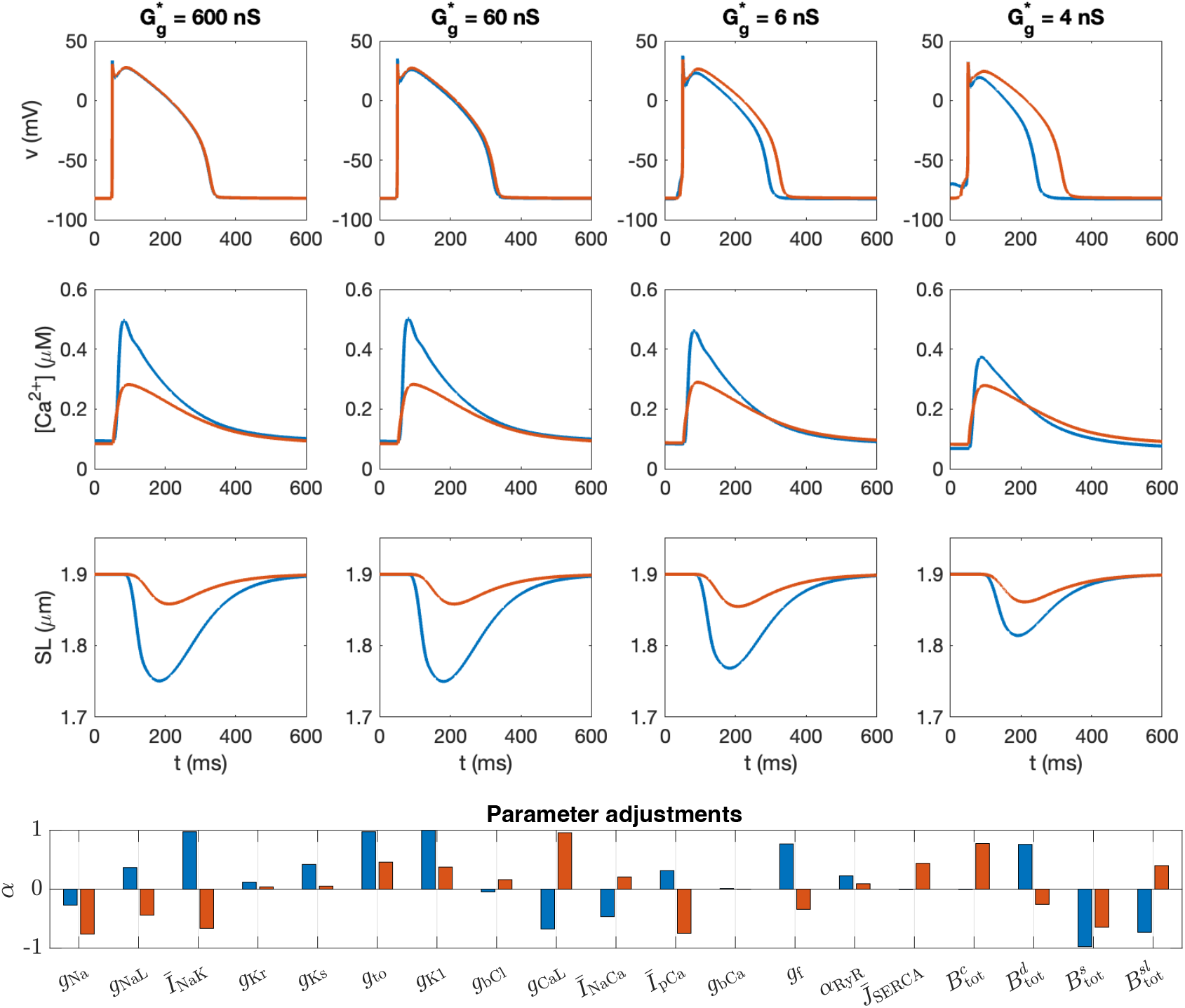
Action potentials, calcium transients and sarcomere length (SL) shortening for two different cells in a collection of 25 *×* 25 cells with 50% variation in cell properties and gap junction coupling (Case 5 in Table 1, Figure 5). We plot the properties of the two cells along the same *x*-values with the largest difference in the action potential duration (APD50) for the 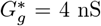 case. The lower panel displays the parameter adjustments, *α*_*i,k*_ (see (7)), corresponding to the two cells.

### 3.5 The repolarization wave is more sensitive to cell perturbations than the depolarization wave

Figure 8 displays the results of simulations similar to those displayed in Figure 4, but where cell variation is represented by perturbation currents that are constant in time instead of by adjustments of ion channel conductances and other model parameters (Case 6 in Table 1). The gap junction coupling is the same between all cells, but we consider four different values of gap junction coupling strength. We observe similar effects as those observed in Figure 4 also in this more simplified example. Moreover, for the medium gap junction conductances of 60 nS and 6 nS, there is considerably more irregularities in the wavefronts during repolarization than during depolarization, indicating that the repolarization wavefront is more sensitive than the depolarization wavefront to this type of cell perturbation.

**Figure 8:**
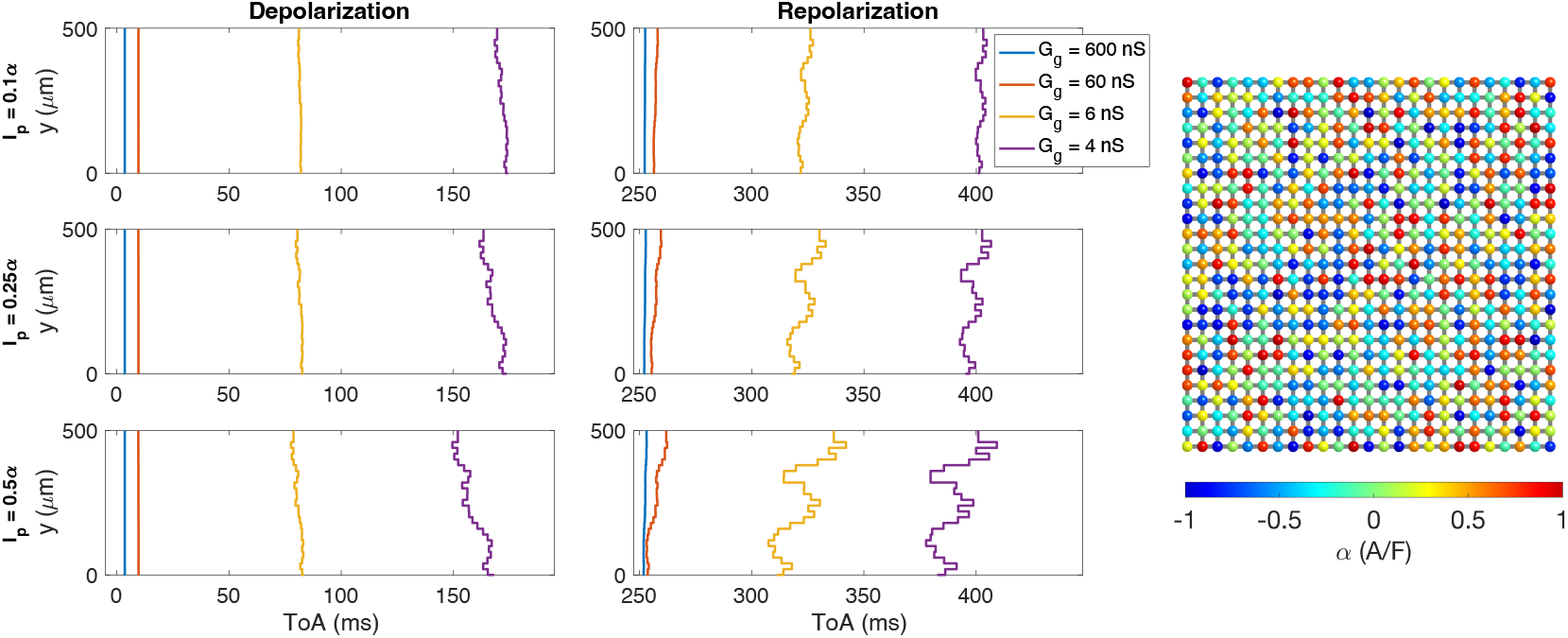
Time of arrival (ToA) for the depolarization and repolarization waves at cell number 13 in the *x*-direction and each of the cells in the *y*-direction in simulations of 25*×* 25 cells with four different gap junction conductances (see legends). The strength of the gap junction coupling and the cell properties are the same for all cells, except that a perturbation current, *I*_*p*_, that is constant in time, is included for each cell (Case 6 in Table 1). This current is given on the form *I*_*p*_ = *rα*, where *α∈* [*−*1 A/F, 1 A/F] varies randomly between cell (see right panel), and *r* is increased for each row. Note that the ToA axis covers different time points in the different plots, but the scaling of the axis is the same in all cases.

### 3.6 Cell heterogeneity and reduced gap junction coupling may form a substrate for EADs

Next, we consider the case of early afterdepolarizations (EADs). For these investigations, we apply the O’Hara-Rudy model with a slow pacing rate (0.25 Hz) and a considerably increased L-type calcium current (*g*_CaL_ multiplied by 4). In addition, we consider a 53% block of the *I*_Kr_ current. This blocking percentage is not strong enough to initiate EADs in the default single cell case, but EADs appear in the default single cell case with 59% block of *I*_Kr_ (in combination with 0.25 Hz pacing and 4 *× g*_CaL_).

#### Variation in gap junction coupling

In Figure 9, we observe that EADs are also not initiated in a collection of cells with variation in the gap junction coupling, but no variation in cell properties (Case 3 in Table 1). The lower panel of the figure illustrates the simulation set up with different degrees of variation in the gap junction coupling. In addition, seven cells along the *x*-direction of the cell collection is marked. The membrane potential of these seven cells are plotted in the upper panels of the figure. We consider three different degrees of gap junction variation (0%, 25% and 50%), as well as five different values for the average gap junction coupling (600 nS, 60 nS, 6 nS, 4 nS, and 2 nS). We observe that for strong cell coupling, the action potentials occur almost synchronously for all the cells, whereas for a weak cell coupling (e.g., 2 nS), a traveling excitation wave is clearly visible with both depolarization and repolarization occurring at different points in time for the different cells. Nevertheless, as noted above, EADs are not initiated in any of the considered cases. This is as expected since the properties of all cells are the same and such that EADs are not initiated in the single cell case.

**Figure 9:**
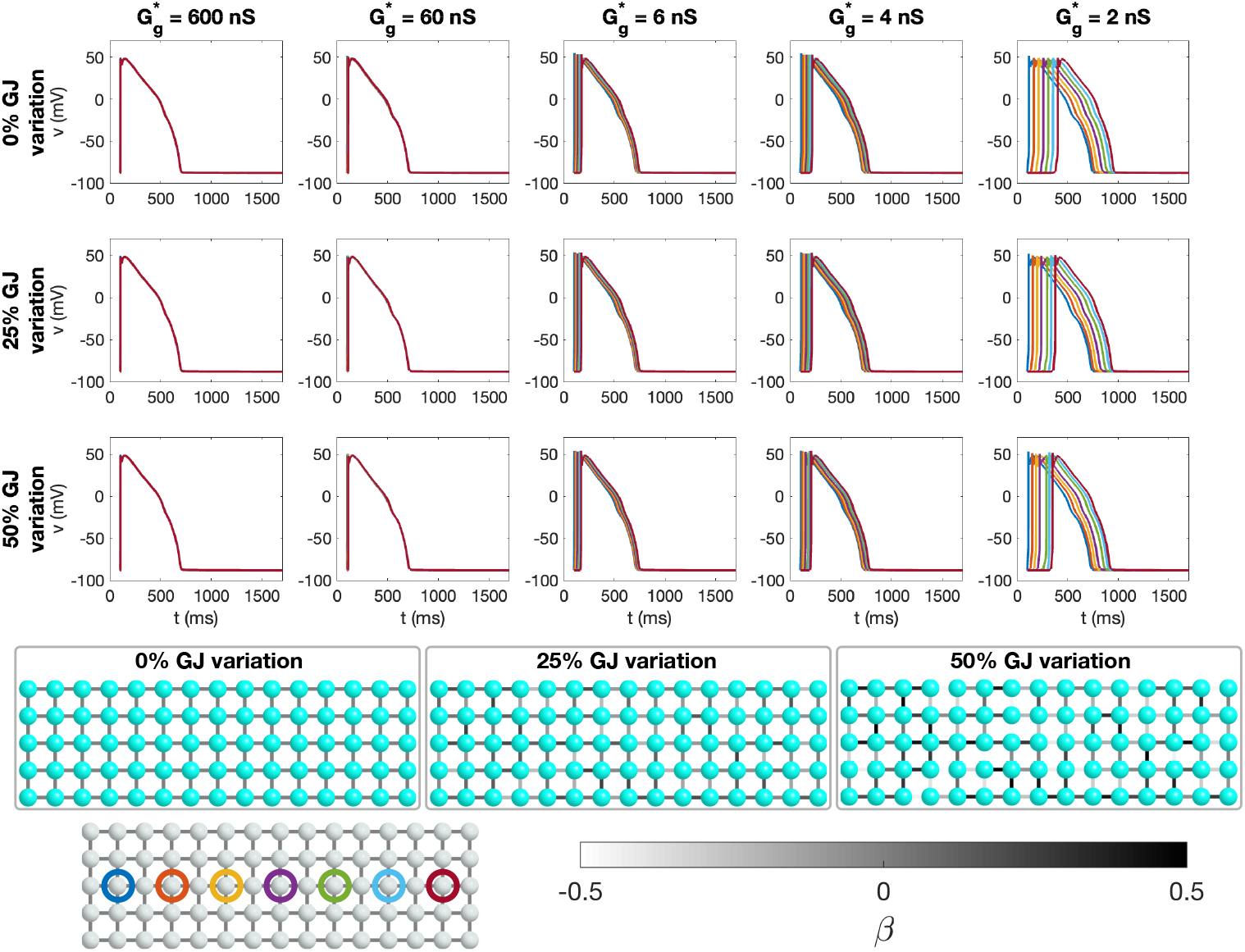
Simulation of a collection of 15*×*5 cells modeled using the O’Hara-Rudy model with 53% *I*_Kr_ block, 4*×g*_CaL_ and 0.25 Hz pacing. We consider five different values of the average gap junction conductance (column titles). The cell properties are the same for all cells. The gap junction conductance is constant in the upper row and varies randomly between cell connections within *±*25% and *±*50% of the average gap junction conductance, respectively, in the center and lower rows (Case 3 in Table 1). We plot the action potential in the cells along a line in the *x*-direction. In the lower left panel, these cells are circled with a color corresponding to the color of the plotted action potentials.

#### Variation in cell properties

In Figure 10, we consider variation in cell properties instead of variations in gap junction coupling (Case 4 in Table 1). The gap junction coupling strength is the same for all cell connections and the parameters of the O’Hara-Rudy model is varied randomly for each cell as explained in Section 2.3.1. The lower figure panel illustrates the variation in cell properties by reporting the APD50 value for each cell if it was considered in isolation and with no *I*_Kr_ block. In the upper panels of the figure, the membrane potential for some of the cells in the cell collection are displayed in the same manner as in Figure 9. In this case, we observe that for a weak cell coupling (2 nS) in combination with 25% or 50% cell variation, EADs are initiated. Furthermore, for the 4 nS cell coupling, EADs are only initiated for a high degree of cell variation (50%). This indicates that a collection of heterogeneous cells that are safe from EADs when the gap junction coupling is normal, may be vulnerable to EADs if the gap junction coupling is sufficiently reduced.

**Figure 10:**
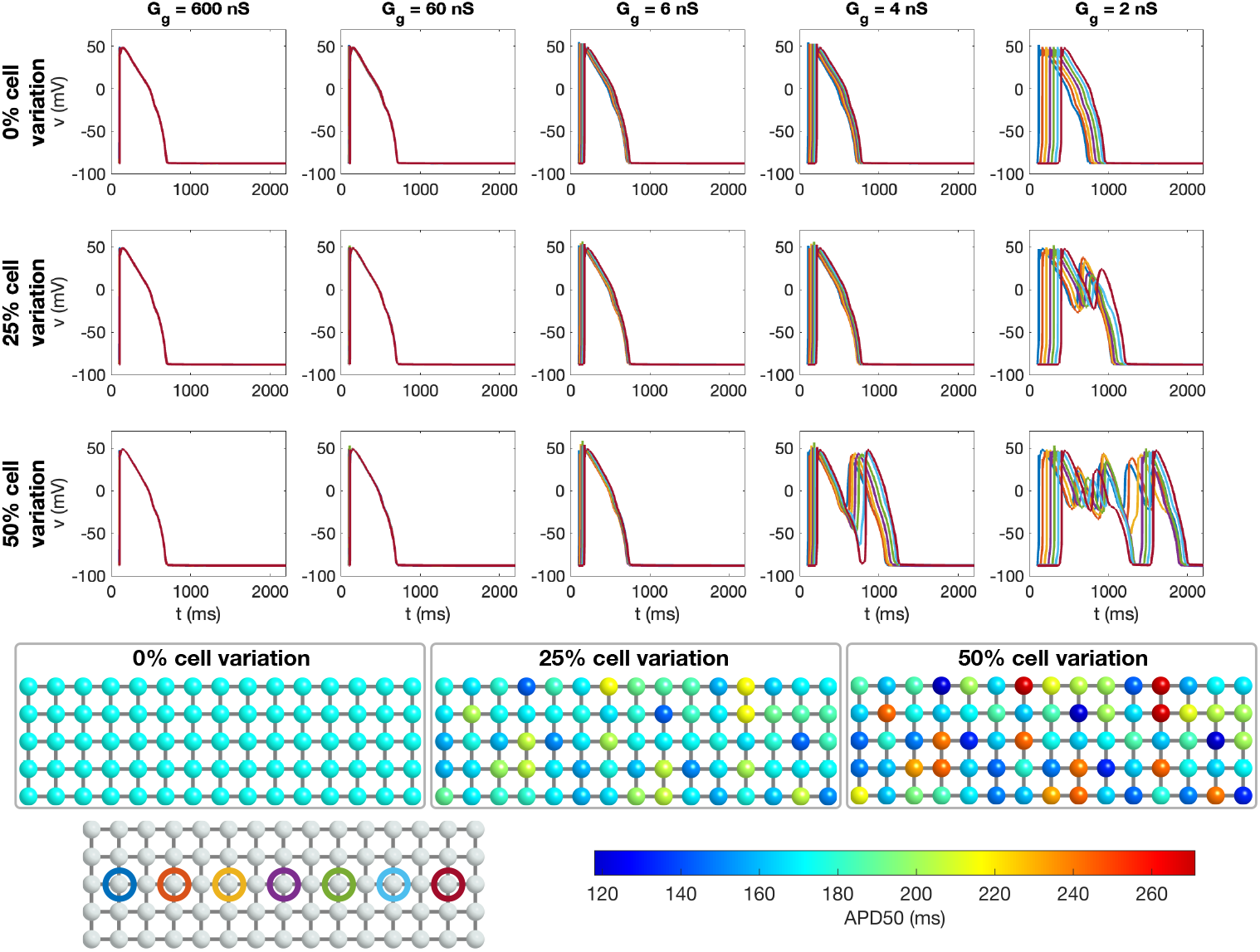
Simulation of a collection of 15*×*5 cells modeled using the O’Hara-Rudy model with 53% *I*_Kr_ block, 4*× g*_CaL_ and 0.25 Hz pacing. We consider five different values of the average gap junction conductance (column titles). The cell properties varies randomly from cell-to-cell (Case 4 in Table 1), and the isolated single cell APDs corresponding to 1 Hz pacing and no parameter adjustments are displayed in the lower panel. The degree of variation of cell properties are increased for each plot row. The gap junction conductance is the same, *G*_*g*_, between all cells. We plot the action potential in the cells along a line in the *x*-direction. In the lower left panel, these cells are circled with a color corresponding to the color of the plotted action potentials.

#### Variation in cell properties and gap junction coupling

Finally, in Figure 11, we consider the case of a collection of cells with variations in both the gap junction coupling and in the cell properties (Case 5 in Table 1). Again, we observe that EADs are initiated for a high degree of cell variation and a weak gap junction coupling.

**Figure 11:**
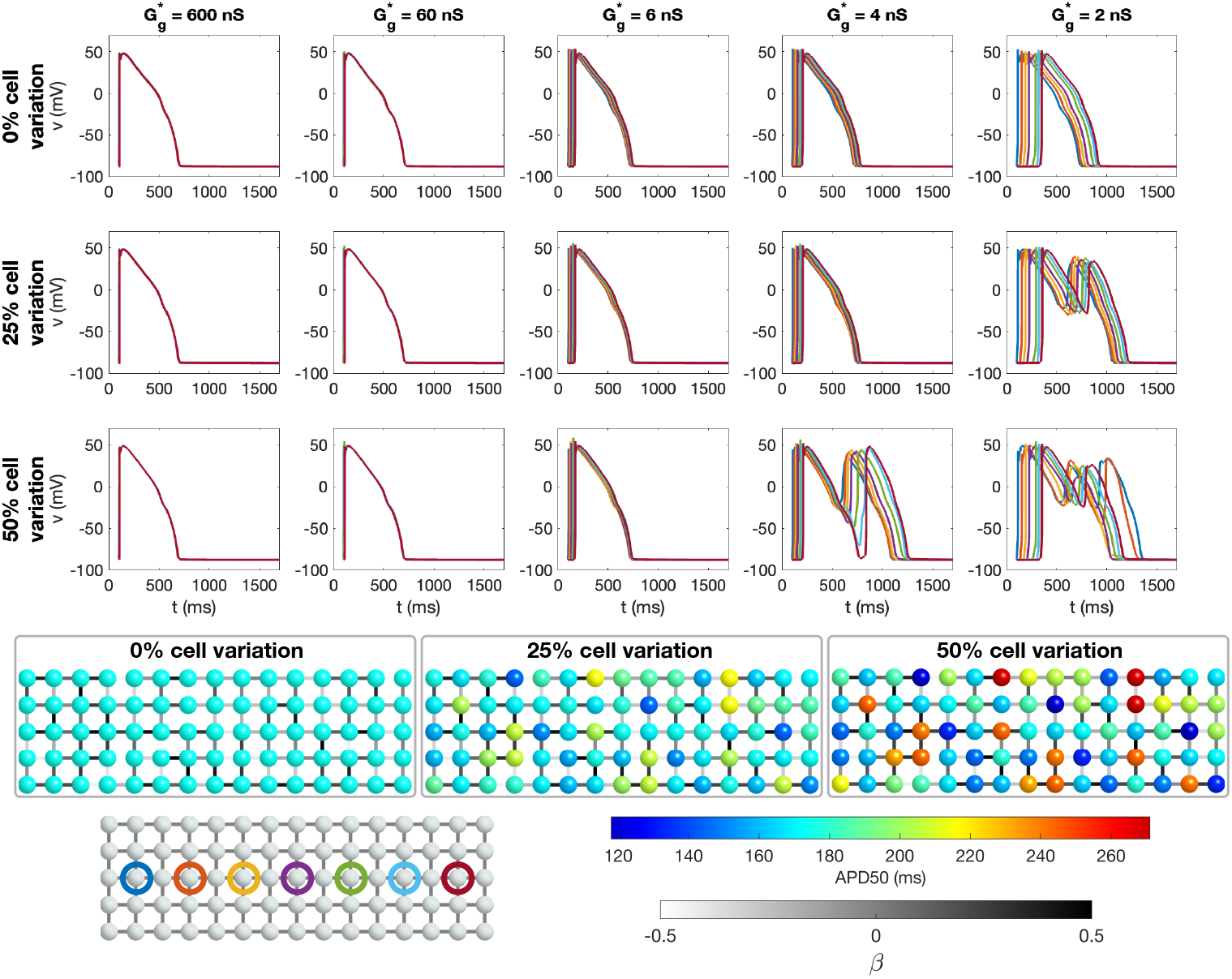
Simulation of a collection of 15*×*5 cells modeled using the O’Hara-Rudy model with 53% *I*_Kr_ block, 4 *×g*_CaL_ and 0.25 Hz pacing. We consider five different values of the average gap junction conductance (column titles). The cell properties varies randomly from cell-to-cell, and the isolated single cell APDs corresponding to 1 Hz pacing and no parameter adjustments are displayed in the lower panel. The degree of variation of cell properties are increased for each plot row. The gap junction conductance varies randomly between cell connections within *±* 50% of the average gap junction conductance, 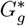. We plot the action potential in the cells along a line in the *x*-direction. In the lower left panel, these cells are circled with a color corresponding to the color of the plotted action potentials.

Figure 12 shows the *I*_CaL_ current associated with the cell marked with blue in the lower part of Figure 11. We consider the case of 50% cell variation and 50% gap junction variation (Case 5 in Table 1). We observe that for strong cell coupling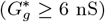, a small increase in the *I*_CaL_ current is present at the end of the action potential, but it is not sufficient for an EAD to be generated. For the weaker cell coupling, however, *I*_CaL_ increases enough to trigger EADs.

**Figure 12:**
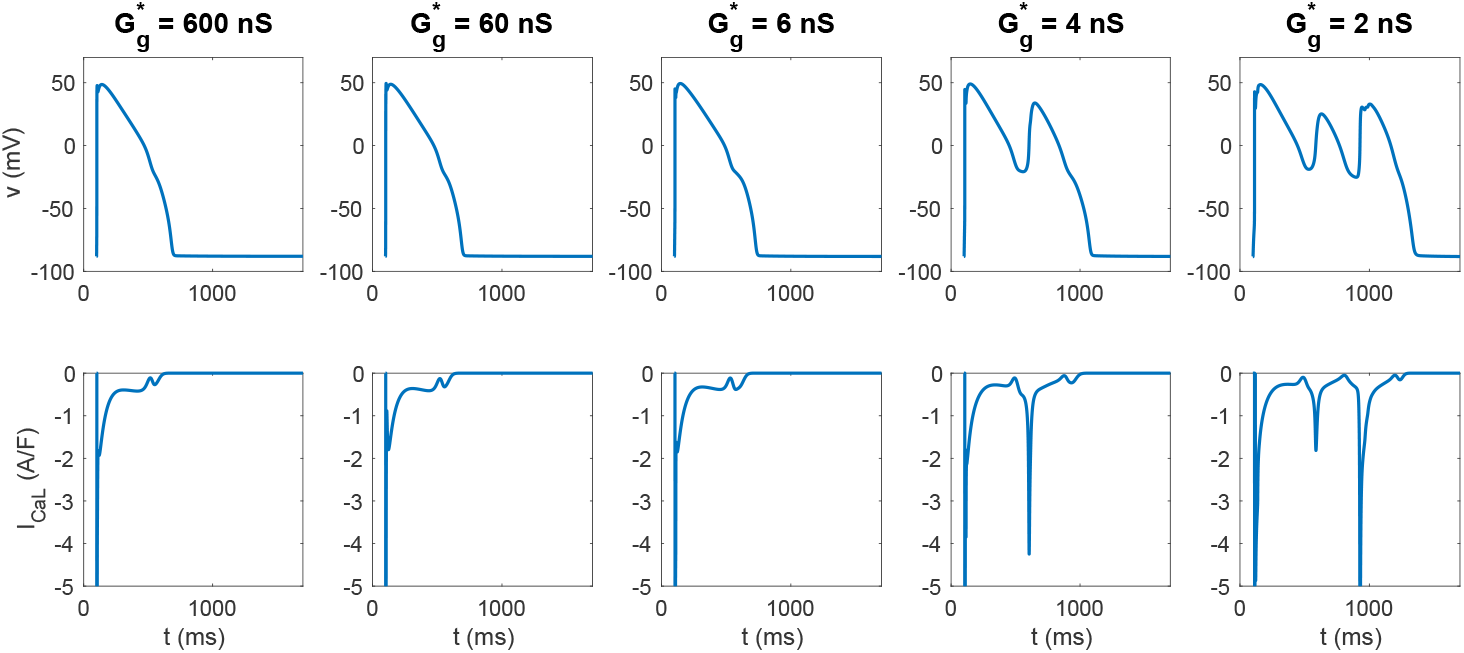
Action potentials and *I*_CaL_ currents in the cell marked with a dark blue circle in the lower panel of Figure 11. We consider the case of 50% cell variation and 50% gap junction variation (Case 5 in Table 1)

## 4 Discussion

### 4.1 Strong gap junction coupling homogenizes cellular heterogeneity

The main observation of this paper is that strong cell-to-cell gap junction coupling significantly improve the electrical conduction of cardiac tissue in the sense that the excitation front becomes synchronized as the coupling strength increases. We observe this in Figure 1, where two cell types with very different properties result in convergent properties as the cell-to-cell coupling is increased. Interestingly, the convergence is weaker at the boundary cells (see Figure 2), which may have implications for border regions such as the border zone of an infarcted region. Additionally, when gap junction coupling is strongly varying, when the cell properties are strongly varying, or when both factors are present (see Figures 3, 4, 5), sufficiently strong cell-to-cell coupling results in converged properties and more or less straight excitation fronts. A similar observation holds in the case of including perturbation currents; strong cell-to-cell coupling leads to convergence and repair of the ragged front present for low cell-to-cell coupling, see Figure 8. This aligns with previous findings on the role of gap junction coupling in stabilizing conduction and reducing arrhythmic susceptibility, see, e.g., [49].

### 4.2 Strong cell-to-cell coupling does not fully synchronize contraction

The pronounced convergence of membrane potential properties across different cells due to increased cell-to-cell coupling is not equally reflected in calcium transients or contraction dynamics, as shown in Figure 7. Unlike the membrane potential, cell-to-cell coupling does not directly influence calcium transients or cellular contraction but rather exerts its effects indirectly through changes in the membrane potential. As a result, there is no direct mechanism enforcing convergence in the contractile machinery across cells, even though the initiation of contraction is clearly synchronized via the membrane potential. This distinction explains the limited extent to which electrical coupling alone can homogenize mechanical responses in cardiac tissue.

### 4.3 Development of EADs depends on weak cell-to-cell coupling

Early afterdepolarizations (EADs) are believed to be a major contributor to serious ventricular arrhythmias, see, e.g., [50, 51]. In the modeling framework applied here, we observe in Figures 9, 10 and 11 that EADs only appear when cell-to-cell coupling is very weak.

This suggests that when gap junction coupling is strong, neighboring cells effectively dampen abnormal depolarizations, reducing the likelihood of EAD development. Furthermore, it suggests that electrotonic couplings stabilize the repolarization wave. Conversely, under weak coupling conditions, individual cells become more isolated, allowing localized afterdepolarizations to persist and potentially trigger arrhythmic events.

EAD formation is closely linked to prolonged action potential duration and disruptions in ionic balance, particularly involving calcium and potassium currents, [52]. In the model, weak cell-to-cell coupling clearly exacerbates the effects of these conditions by reducing the stabilizing influence of neighboring cells on repolarization. This may be particularly relevant in diseased states, where structural remodeling and fibrosis impair electrical connectivity. hERG (*I*_Kr_) block is a well-known but unintended side effect of certain drug compounds, and its risks may be significantly heightened in patients with reduced cell-to-cell coupling, [53].

### 4.4 Improved gap junction coupling as a therapeutic option

We have seen that strengthening the gap junction coupling strongly homogenizes the membrane potential of cardiomyocytes with highly variable membrane properties. It synchronizes the conduction wave and reduces wavefront irregularities. Furthermore, it is well known that increased cell coupling also increases the conduction velocity, which may reduce the probability of dangerous re-entry waves, see, e.g., [54, 55, 56, 25]. Therefore, increasing the gap junction coupling appears to be a promising candidate for therapeutic intervention. However, no such drug has yet reached the market. One challenge is that enhancing gap junction coupling can also facilitate fibroblast-cardiomyocyte interactions, potentially promoting arrhythmogenic effects in certain conditions, [57]. Additionally, the role of gap junctions in cancer development is complex, [58]. While connexins have been implicated in tumor suppression in some contexts, increased intercellular coupling may also facilitate cancer progression under certain conditions. A more direct approach to modifying intercellular coupling is the overexpression of gap junction proteins via gene therapy; increased Cx43 expression has been shown to improve conduction velocity and reduce atrial fibrillation susceptibility, [59].

## 5 Conclusion

We have demonstrated how gap junction coupling influences the propagation of excitation waves in cardiac tissue, particularly in the presence of substantial cellular heterogeneity. When coupling is strong, the effect of variability in membrane properties is largely suppressed, resulting in synchronized and smooth conduction. However, as gap junction conductance decreases, the effects of cell-to-cell differences become more pronounced, leading to irregular wavefronts and increased susceptibility to early afterdepolarizations. Notably, while electrical coupling significantly homogenizes action potential dynamics, it does not fully synchronize calcium transients or mechanical contraction. Our findings confirm that reduced gap junction coupling, as observed in fibrotic regions, contributes to conduction instability and may facilitate arrhythmogenesis. The results underline the importance of considering cellular interactions in studies of cardiac electrophysiology, rather than relying solely on single-cell models.

